# Proteomic characterisation of Japanese quail’s unique seminal foam

**DOI:** 10.1101/2025.04.23.650219

**Authors:** Chloe Mason, Martin Garlovsky, Oscar Vedder, Trong Khoa Pham, Rachel George, Barbara Tschirren, Nicola Hemmings

**Author notes:** **Corresponding Author: Chloe Mason**, University of Sheffield, Alfred Denny Building, Western Bank, Sheffield, S10 2TN.

## Abstract

Male Japanese quail (*Coturnix japonica*) produce a unique proctodeal foam upon ejaculation which is thought to enhance fertilisation success. However, the precise function and proteomic characterisation of this foam remain largely unexplored. Using high throughput LC-MS/MS proteomics analysis, we characterised the proteome of the quail seminal foam for the first time. Our analysis confidently identified 224 proteins in the foam using 32 pooled samples from 96 males. 96.4% of proteins had orthologs in the closely related (diverged 35 million years ago) chicken (*Gallus gallus*) which generated 40 Gene Ontology terms. Further interrogation revealed proteins involved in sperm motility, maturation and DNA protection, suggesting the foam plays a role in fertilisation success. Proteins linked to immune defence and inflammation modulation were identified, indicating the foam may protect sperm from antigens and reduce female immunogenic responses. This work advances our understanding of the function and evolution of Japanese quail unique seminal foam and directs future experiments to investigate the role of key seminal foam proteins in post-copulatory sexual selection.

**Significance of the Study:** Seminal fluid plays an important role in fertilisation success, yet the molecular components of seminal fluid and their function remain poorly understood in most species. This study provides the first comprehensive proteomic characterisation of a unique seminal foam produced by the Japanese quail (*Coturnix japonica*), revealing its potential mechanistic function in sperm regulation, sperm motility, immune protection, and fertilisation success. By identifying key proteins associated with energy production, membrane fluidity, and sperm maturation, our findings suggest the foam may influence post-ejaculatory sperm function. Additionally, the identification of immunity proteins indicates a role in sperm protection and modulation of female immune responses. This study advances our understanding of avian reproductive biology and opens new avenues for investigating the specific role of seminal foam proteins in fertilisation dynamics and post-copulatory sexual selection.

## 1 Introduction

Upon ejaculation, males transfer not only sperm but also seminal fluid, which has a range of effects on sperm and the female environment it operates in. Seminal fluid has components necessary for sperm viability, storage, maturation, and ultimately fertilisation success. In particular, seminal fluid proteins have been shown to play a vital role in mediating sperm performance and modulating female reproductive physiology [1–4] and exhibits rapid rates of evolution across species [5–7].

Male quail of the genus *Coturnix* are unique among birds for producing a white foam upon ejaculation, in addition to seminal fluid, which is secreted from a specialised gland known as the ‘proctodeal gland’ or ‘cloacal gland’ [8]. The foam is a viscous glycomucoprotein aerated by male cloacal muscle contractions [9], as well as interactions with carbon dioxide and hydrogen produced by cloacal bacteria [10]. As in chickens (*Gallus gallus*) and turkeys (*Meleagris gallopavo*), seminal fluid is derived from the testis, epididymis, and ductus deferens in Japanese quail (*Coturnix japonica*) [11], but foam is produced separately and not mixed with sperm or seminal fluid until deposited into the female cloaca during copulation [8]. The foam’s function is not fully understood; unlike seminal fluid, it is not required for fertilisation [12, 13] but may aid sperm function in the oviduct and improve fertilisation success [14, 15].

Male proctodeal gland size is a predictor of fertilisation success, and natural copulations with foam have higher fertilisation success compared to without foam, or with artificially placed foam [16–18]. Foam may extend the female’s fertile period, increasing the likelihood of successful fertilisation [15, 17] or improve sperm motility. Foam significantly prolongs sperm motility and increases sperm velocity in vitro, suggesting its components supply energy to sperm [14, 19]. Lactate dehydrogenase is a protein found at high levels in quail seminal plasma [20], and lactate found in foam may act as an energy source for sperm transport [14] as it does in the seminal fluid of mammals [21, 22]. Foam also disaggregates sperm clumps, leading to more vigorous motility, possibly due to non-protein components [14]. Most studies investigating the effect of foam on male fertilisation success have employed a foam removal manipulation technique confounded by invasive surgery likely to interfere with copulation. However, Finseth et al. [23] showed that non-invasive foam removal in natural mating scenarios reduces male fertilisation success under sperm competition from rival males. The function of foam during sperm competition, possibly mediated through positive effects on own sperm at a cost to a rival’s fertility, suggests it evolved under sexual selection [23].

The broader protein composition and function of quail foam have not yet been studied but could offer an interesting comparison to seminal fluid proteins that are differentially produced but play a similar role in other Galliformes [24–26]. Here, we use high-throughput proteomics using liquid chromatography-mass spectrometry (LC-MS) to characterise the protein composition of the unique seminal foam of the Japanese quail for the first time, offering an improved inference on its evolution and function.

## 2 Materials and Methods

### 2.1 Collection of Seminal Foam

We collected foam samples from two cohorts (in 2018 and 2019) of a population of captive Japanese quail (*Coturnix japonica*) housed at the Institute of Avian Research, Wilhelmshaven, Germany, consisting of 17 and 79 adult males, respectively. All birds were kept under licence of the Veterinäramt JadeWeser (permit number 42508 03122020). Foam was collected by gently squeezing the proctodeal gland and stored at −80°C until analysis. Foam samples were defrosted at room temperature for 12 hours and pooled within cohorts (to ensure sufficient material for mass spectrometry analysis) to give a total of 32 replicate samples (8 replicates from 2018 and 24 replicates from 2019). We combined data from both cohorts to provide an overall characterisation of the foam proteome.

### 2.2 Sample Preparation

To isolate the foam proteome, we added four times the sample volume of ice-cold acetone (−20°C) to the sample, vortexed and incubated it for 4 hours at 20°C, and then centrifuged it for 10 minutes at 13,000rpm at 4°C. Supernatant was discarded, and the protein pellet was left at room temperature until the acetone evaporated completely.

Protein pellet (∼25µg) was then resuspended in 25µL protein lysis buffer (5% SDS (Sigma-Aldrich) and 100mM TEAB (ThermoFisher), pH 8.5), then reduced by 20mM DTT (ThermoFisher) using a thermoshaker at 95°C, 800rpm for 10 minutes. After cooling for 5 minutes at room temperature, proteins were alkylated using 40mM 2-iodoacetamide (Sigma-Aldrich) using a thermoshaker at room temperature, 800rpm for 30 minutes in the dark. Proteins were then digested using a suspension trapping (S-Trap) technique according to manufacturer’s protocol (ProtiFi). Briefly, alkylated proteins were acidified by adding 2.5µL 12% phosphoric acid, followed by 365µL S-trap binding buffer (90% aqueous methanol, 0.1M TEAB, pH 7.1). Samples were then transferred to S-Trap columns (Protifi) gently and centrifuged for 60 seconds at 4000xg to trap the denatured proteins.

Trapped proteins were washed 5 times with 150µL binding buffer, centrifuging between each addition, then transferred to clean 1.5ml Eppendorf tubes for protein digestion. To digest proteins into peptides, 25µL MS grade Trypsin (ThermoFisher) in 50mM TEAB buffer (concentration 0.1µg/µL) was added to each S-Trap and incubated at 47°C for 1.5 hours without shaking. Peptides were eluted using a series of solvents: 40µL 50mM TEAB, 40µL 0.2% aqueous formic acid (ThermoFisher), 40µL 50% ACN in 0.2% formic acid, and 40µL 80% ACN in 0.2% formic acid. Samples were centrifuged at 4000xg for 60 seconds between solvent additions. Eluted peptides were collected and dried in a vacuum concentrator (Eppendorf) for 2 hours before being reconstituted in 60µL 0.5% formic acid and 4µL withdrawn for MS analysis.

### 2.3 LC-MS/MS Analysis

We performed sample processing separately for each cohort. All MS proteomics analyses were performed at the bioMICS Mass Spectrometry Facility (https://www.sheffield.ac.uk/mass-spectrometry/biomics), University of Sheffield, on an Orbitrap Exploris^™^E480 mass spectrometer (ThermoFisher) equipped with a nanospray source, coupled to a Vanquish HPLC System (ThermoFisher). Peptides were desalted online using a Nano-Trap Column (75μm I.D.X 20mm; ThermoFisher) and then separated using an EASY-Spray^TM^ column (50cm×50μm I.D., PepMap C18, 2μm particles, 10Å pore size; ThermoFisher). We used a 100-min gradient, starting from 3% to 20% buffer B (0.5% formic acid in 80% ACN) for 68 minutes, followed by a ramp-up to 35% buffer B for 23 minutes, then to 99% buffer B for 1 minute, and maintained at 99% buffer B for 9 minutes. The MS was operated in positive mode with a cycle of 1 MS acquired at a resolution of 120,000, at m/z of 400. The top 20 most abundant multiply charged (2^+^ and higher) ions in a given chromatographic window were subjected to MS/MS fragmentation in the linear ion trap. The scan range (m/z) was 375–1200, with a normalised AGC target of 300%, microscan 1, an FTMS target value of 1e4, and a resolution of 15,000.

### 2.4 Protein Identification and Bioinformatic Analysis

All MS data were analysed using MaxQuant (v.1.6.10.43) and searched against the *Coturnix japonica* protein database consisting of 27,875 proteins (UniProt proteome ID: UP000694412) with the following search parameters: trypsin/P (2 missed cleavages) as the enzyme, methionine oxidation and N-terminal protein acetylation as variable modifications, and cysteine carbamidomethylation as a fixed modification. FDR of 0.01 was used for both peptide and protein identification.

The MaxQuant output was loaded into Perseus (v.1.5.6.0) and all label-free quantification (LFQ) intensities were set as main columns. The matrix was filtered to remove all proteins that were contaminants and reverse sequences. Only proteins identified in at least 3 replicates within a cohort were included in the final protein list. LFQ intensities were then transformed using the log_2_(x) function. The quality of replicates were examined using Pearson’s correlation analysis.

### 2.5 GO Analysis

As the *C. japonica* genome has not been fully annotated, we identified chicken (*Gallus gallus domesticus*) orthologs of quail foam proteins using OrthoFinder (v.3.0) [27] and the chicken reference proteome (UniProt Proteome ID: UP000000539). A Chi-square test performed in R (v.4.4.2) [28] was used to compare the proportion of foam proteins with chicken orthologs to the proportion of all Japanese quail proteins with chicken orthologs. The list of chicken orthologs and their relative abundance from both cohorts were combined and any replicates were removed. We assessed the overlap between the quail foam proteome, the seminal fluid proteome of the red junglefowl [24] and domestic chicken [25], and the chicken spermatozoa proteome [25].

GO analysis was performed on the chicken orthologs using the website version of DAVID [29, 30]. The proteins list (foreground = 449) was uploaded to DAVID (https://davidbioinformatics.nih.gov/tools.jsp) and the *G. gallus* official gene list was used as the background. Outputs for all three direct GO categories (biological process, cellular components, and molecular functions) were downloaded including the associated statistical values, as well as the output table from the Functional Annotation Clustering Tool that clusters redundant annotation terms to identify biological themes associated with the proteome. Figures were created using R (v.4.4.2) [28].

## 3 Results

We identified 608 proteins in Japanese quail foam across the two cohorts (Table S1), of which 224 were identified in at least 3 replicates in a single cohort (36.8%) (Table S2). The relative protein abundances were measured using LFQ, and their abundances were strongly correlated between replicates (mean Pearson’s correlation coefficient = 0.85, range = 0.55 – 0.99). See Table 1 for the 20 most abundant proteins in the foam.

**Table 1:**
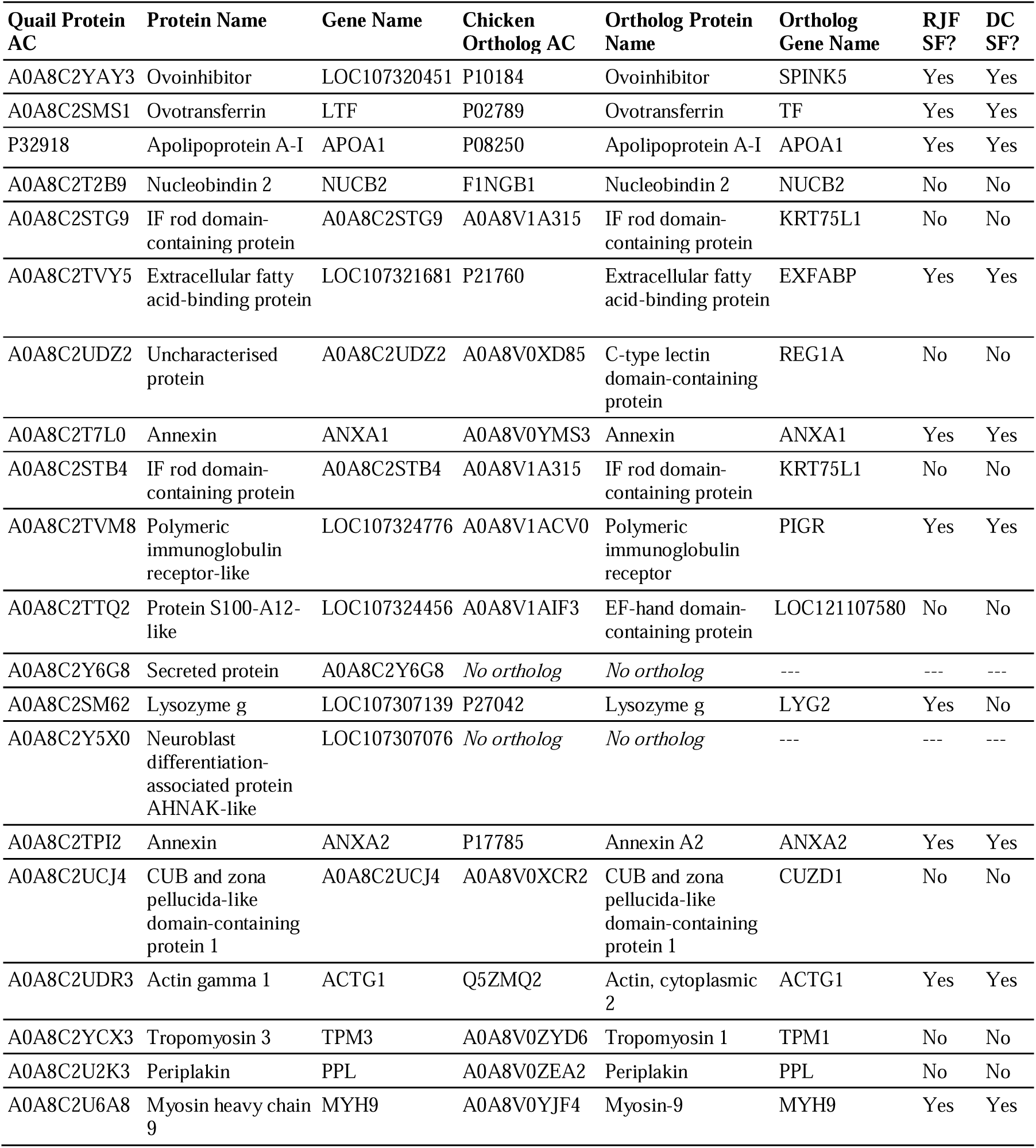
The 20 most abundant proteins in Japanese quail (*Coturnix japonica*) seminal foam by LFQ (rank ordered by abundance), and their chicken (*Gallus gallus domesticus*) orthologs. The chicken orthologs were compared to the proteins found in the red junglefowl seminal fluid (RJF SF) and the domestic chicken seminal fluid (DC SF).

**Table 2:**
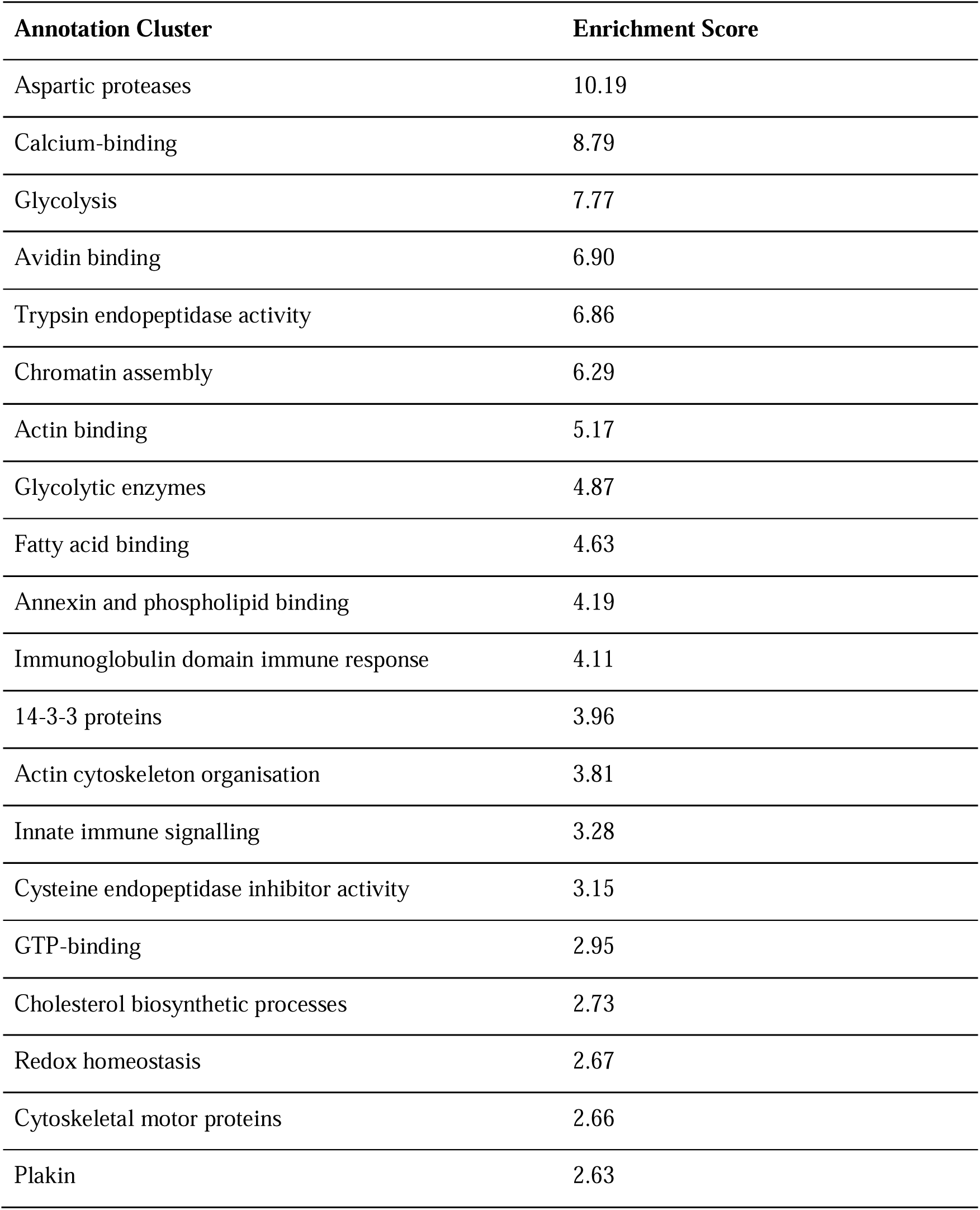
Functional annotation clusters that are significantly enriched in Japanese quail (*Coturnix japonica*) seminal foam proteome. Only clusters with an enrichment score >2.00 and terms with Benjamini-Hochberg corrected *p*-value <0.01 are shown. See Table S5 for a full list of annotation clusters, associated terms and *p*-values.

### 3.1 Chicken Orthologs

Genes encoding foam proteins are highly conserved, with 96.4% of proteins (216/224) having chicken orthologs (Table S2) compared to 82.0% of all Japanese quail proteins (22,858/27,875; χ^2^= 31.17, df = 1, *p* < 0.001). More than one chicken ortholog was found for 27.7% (62/224) of foam proteins (Fig. 1), and there were 449 chicken proteins orthologs in total encoded by 266 chicken ortholog genes.

**Figure 1:**
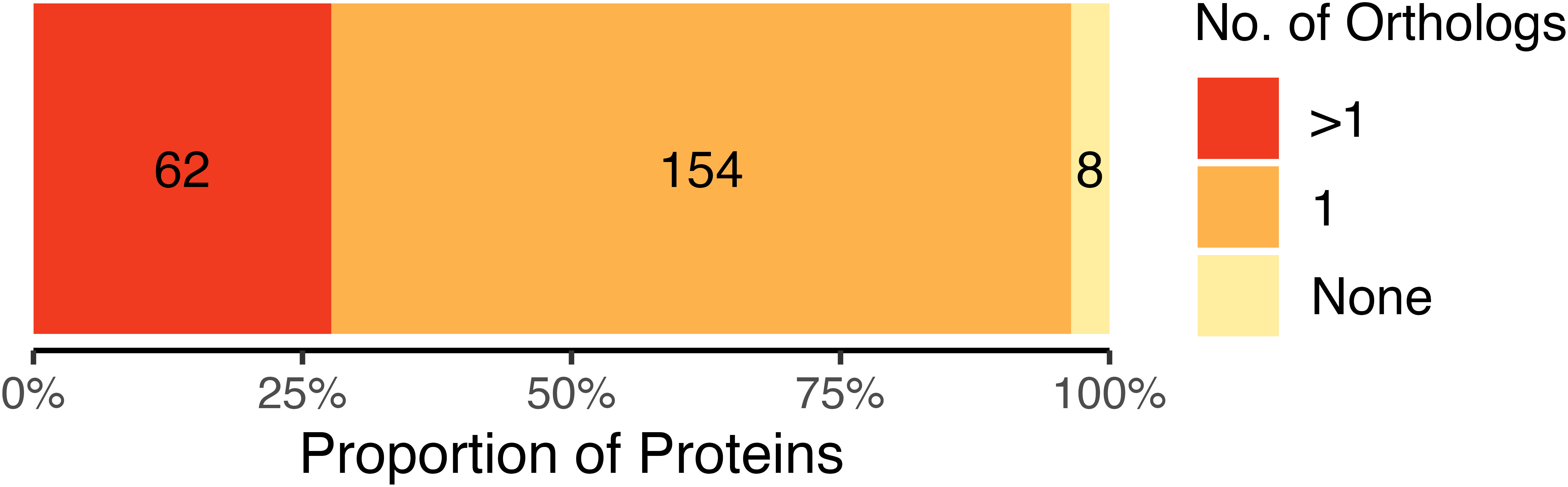
The proportion of Japanese quail (*Coturnix japonica*) seminal foam proteins that have more than one (red), one (orange), or no (yellow) chicken (*Gallus gallus domesticus*) orthologs.

Eight foam proteins did not have chicken orthologs (Table S3) and their functions were inferred using the UniProt database [31]. Four are predicted (with no evidence of existence) to be neuroblast differentiation-associated protein AHNAK-like (3 of which are protein isoforms) and associated with the nucleus and regulation of RNA splicing. Two predicted Ig-like domain-containing proteins may be associated with an immunoglobulin-mediated immune response. Aldo-keto reductase family 1 member B1 (aldose reductase) was present in foam, and this protein’s existence is inferred from homology, meaning its existence is probable. Finally, a predicted secreted protein associated with the transmembrane helix was present in foam, but no further information on this protein’s function is known.

Comparing proteomes, 7.6% (34/449) and 14.7% (66/449) of chicken orthologs identified in quail foam were also found in domestic chicken and red junglefowl seminal fluid, respectively, and 34.1% (153/449) and 39.6% (178/449) of foam proteins had isoforms found in chicken and red junglefowl seminal fluid, respectively (Fig. 2; Table S4). Comparing genomes, 31.6% (84/266) and 40.2% (107/266) of genes encoding quail foam proteins were found to encode seminal fluid proteins in chicken and red junglefowl seminal fluid, respectively (Table S4). 231 quail foam proteins (141 genes) were not found in either chicken or red junglefowl seminal fluid. GO analysis of proteins unique to quail foam revealed a significant association with proteolysis (Benjamini-Hochberg corrected *p*-value = 4.5E-6, FDR = 4.5E-6).

**Figure 2:**
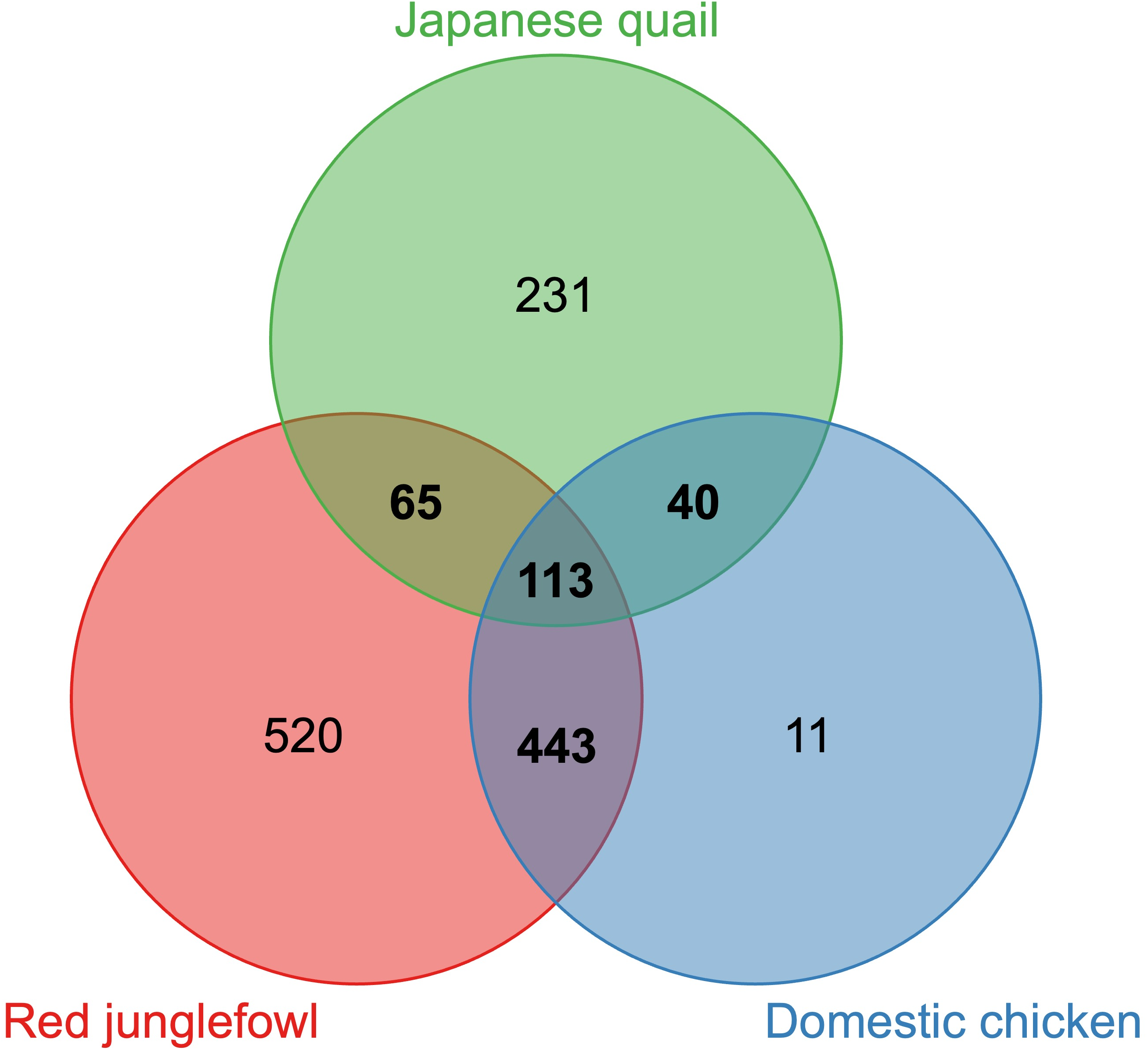
Venn diagram indicating the size and overlap between the seminal foam proteome of the Japanese quail (*Coturnix japonica*) and the seminal fluid proteomes of the red junglefowl (*Gallus gallus*) and domestic chicken (*Gallus gallus domesticus*).

There was evidence that proteins found in chicken sperm are also present in quail foam, with 29.2% of foam protein’s chicken orthologs (131/449) having isoforms identified in chicken spermatozoa (Table S4). Of these, 48 were not found in chicken seminal fluid.

### 3.2 GO Analysis

GO analysis of chicken orthologs identified 7 biological process terms associated with foam (Fig. 3a), including glycolytic process (3.63% of foam proteins; Benjamini-Hochberg corrected *p*-value = 4.52-9, FDR = 4.45E-9), nucleosome assembly (Benjamini-Hochberg corrected *p*-value = 8.35E-6, FDR = 8.22E-6), immune response (Benjamini-Hochberg corrected *p*-value = 1.2E-5, FDR = 1.18E-5), gluconeogenesis (Benjamini-Hochberg corrected *p*-value = 1.2E-5, FDR = 1.18E-5), proteolysis (Benjamini-Hochberg corrected *p*-value = 3.12E-5, FDR = 3.07E-5), fatty acid transport (Benjamini-Hochberg corrected *p*-value = 2.78E-4, FDR = 2.74E-4), and antibacterial humoral response (Benjamini-Hochberg corrected *p*-value = 3.75E-3, FDR = 3.69E-3). Eighteen molecular functions (Fig. 3b) and 15 cellular components (Fig. 3c) were significantly enriched in quail foam (see Table S5 for all GO terms and associated *p*-values).

**Figure 3:**
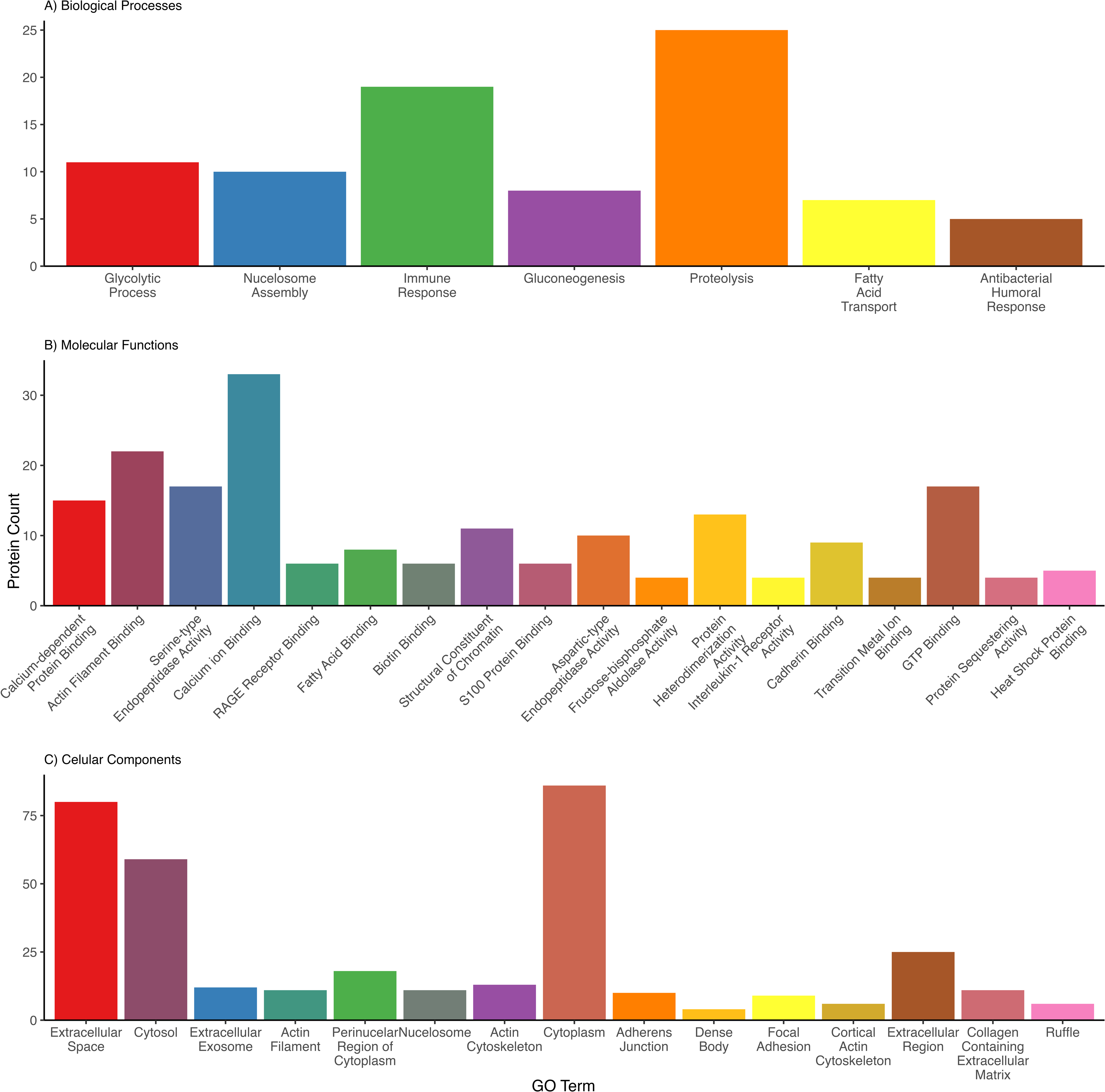
**GO analysis of the Japanese quail (*Coturnix japonica*) seminal foam proteome**: A) 7 biological processes, B) 18 molecular functions, and C) 15 cellular components significantly enriched in the Japanese quail foam proteome, as identified by DAVID. Only functional enrichment groups with Benjamini-Hochberg corrected *p*-values <0.01 and passing a 1% false-discovery rate threshold are shown.

The Functional Annotation Clustering tool in DAVID [29, 30] identified 40 clusters, 20 of which were significantly enriched in the foam proteome (Table 3). Enrichment scores indicated how significantly a cluster is overrepresented in the foam proteome compared to the red junglefowl background proteome (see Table S6 for all annotation clusters, associated terms and *p*-values). As a relatively high number of proteins were associated with proteolysis (Fig. 2a), these proteins were identified individually and assigned to their appropriate protease class (Table S7).

## 4 Discussion

We characterised, for the first time, the proteome of a unique reproductive foam produced by male Japanese quail, identifying a total of 224 proteins. We show over 96% of these proteins have orthologs in the closely related chicken, and at least 48% have been identified in chicken seminal fluid, although this figure may be more with updated proteomic technologies. Previous research has found foam gland genes evolved under strong selective constraint [32], which contrasts with the rapid divergence of testis-derived seminal fluid genes across taxa [5–7].

We identified proteins in foam that are also found in chicken spermatozoa, supporting the idea that foam plays a role in sperm function (29.2% of foam proteins, which is higher than the 9.6% of the red junglefowl seminal fluid proteins) [24, 25]. Unlike chicken seminal fluid, quail foam is produced and stored separately from sperm, and in our study was collected prior to ejaculation, but we cannot rule out the possibility of minor contamination from residual sperm from previous ejaculations in the cloaca.

### 4.1 Role of Foam Proteins in Sperm Regulation and Maturation

Japanese quail have a high turnover of sperm [33] which limits sperm maturation and storage time in the epididymis (a tubular storage/transport organ attached to the testes) [34]. Conversely, female quail store sperm for 8-10 days [35], so maturation may be completed once sperm are transferred to the female. Foam proteins may therefore function during sperm maturation in the female reproductive tract, influencing fertilisation outcomes post-ejaculation. Extracellular vesicles (exosomes) in foam may deliver molecular components to sperm that facilitate maturation [36, 37]. We found proteins associated with extracellular space (GO Term: 0005615) and extracellular exosomes (GO Term: 0070062), including three families of exosome markers that were enriched annotation clusters: (a) GTP-binding regulators, which regulate vesicle formation, trafficking and membrane fusion [38]; (b) annexin proteins, which are membrane scaffolding proteins that regulate vesicle formation [39]and (c) members of the 14-3-3 protein family, which regulate vesicle targeting through cytoskeleton interactions [40], suggesting exosome-like vesicles are abundant in quail foam.

Several annexins identified in foam may bind to phospholipids to stabilise sperm cell membranes [41]. Annexins A1 and A2 were highly abundant in foam, and annexins A7, A8-like protein 1, A11, and A5 were also present. Annexins bind to phospholipid cell membranes in a calcium-dependant manner and have been linked to calcium ion-regulated exocytosis [42] and endocytosis [43], both which may contribute to repair and stabilisation of sperm membranes [44, 45]. Annexin-phospholipid binding also initiates signalling pathways that regulate key cellular processes [41], such as apoptosis [46] and mitosis [47].

Proteolysis (GO Term: 0006508) may be important for regulating sperm maturation and enhancing motility in the female [48, 49], and several proteases (proteolysis enzymes) identified in foam facilitate these functions. Cathepsin D, an aspartic protease, has been shown to function during sperm maturation in humans [50] and mice [51], whilst trypsin, a serine protease, promotes sperm capacitation and activates flagellar activity in the oviduct of insects [52–54]. Several proteases identified in foam may also induce semen liquefaction (where highly viscous semen becomes watery), which is crucial for motility and transport [55]. Trypsin and chymotrypsin, serine proteases in foam, induce liquefaction in vitro in human [56] and spider monkey (*Ateles geoffroyi*) semen [57]. Similarly, metallopeptidases and carboxypeptidases participate in semen liquefaction by remodelling the extracellular matrix [58, 59] and enhancing sperm morphology and viability [60]. Japanese quail foam is known to disaggregate sperm [14], and proteolysis was associated with foam proteins not found in chicken seminal fluid, suggesting this may be a specialised function. Proteases are also known to regulate fertility. In human semen, MMP7 influences apoptosis in the female reproductive tract and is associated with increased fertility [58, 61]. Carboxypeptidase modulates reproductive hormones [62], whilst serine proteases stimulate oviposition and seminal fluid protein transport in insects [63, 64]. Proteolysis is tightly regulated by protease inhibitors to prevent premature activation of pathways or tissue damage [49] and accordingly, we found endopeptidase inhibitors enriched in foam. Despite proteases being common components of accessory gland secretions across species [49], with positive effects on fertilisation [63, 65], little is known about their mechanistic function within the female reproductive tract [49].

Through their role in nucleosome assembly (Go Term: 0006334), several foam proteins may protect sperm DNA. During avian spermatogenesis, the sperm genome compactly transforms into chromatin (a complex of DNA and proteins) that fits inside the sperm head nucleus [66]. Histones are almost completely replaced by protamines in chromosomes, resulting in highly condensed and transcriptionally inactivated chromatin [67, 68]. However, gene transcription and translation may be important for full functionality of ejaculated sperm [69], and there is evidence that the sperm RNA plays a role in supporting embryonic development [70] and embryonic gene expression [71]. Sperm chromatin assembly abnormalities can reduce male fertilisation ability and cause problems during embryonic development [72]. Several histones (H4, H2B and H1.10) identified in foam may originate from minor sperm contamination, as mentioned above, or from cells in the proctodeal gland.

### 4.2 Role of Foam Proteins in Sperm Motility

Across taxa, glycolysis occurs in both seminal fluid and the sperm midpiece, producing ATP from sugars to fuel sperm motility [73]. Glycolysis (GO Term: 0006096) and associated enzymes were enriched in the foam proteome, several of which are also found in chicken and turkey seminal fluid [24, 25] suggesting that foam aids in sperm energy production. Similarly, gluconeogenesis (GO Term: 0006094) and associated enzymes were found in foam, which maintain semen sugar levels produced form noncarbohydrate precursors such as pyruvate and lactic acid [74]. Glycolysis and gluconeogenesis enzymes found in bird and mammalian seminal fluid were originally hypothesised to originate from sperm degradation during epididymal transit or ejaculation [75], but our data suggest in quail these enzymes originate from foam.

We found strong evidence that foam proteins are involved in fatty acid binding (GO Term: 0005504) and transport (Go Term: 0015908), including an extracellular fatty acid-binding protein which was the sixth most abundant protein in foam. Fatty acids esterified to phospholipids form part of the sperm membrane and determine sperm fluidity, flexibility and receptor function [76–78], essential for sperm movement, membrane fusion during fertilisation, cell adhesion, and signal transduction during spermatogenesis [79, 80]. Albumin, which is found in foam and is the most conserved protein in red junglefowl and chicken seminal fluid [81], stimulates sperm motility by binding to phospholipids, reducing the cholesterol/phospholipid ratio in the sperm plasma membrane [82] and inducing capacitation Abnormal lipid homeostasis can cause spermatogenic dysfunction and consequent infertility [83].

The cholesterol-to-phospholipid ratio is critical for normal sperm morphology, sperm motility and fertilisation ability [84–87]. Apolipoprotein A-1 (apoA-1), the third most abundant protein in foam, may be useful during cholesterol biosynthetic by binding to high-density lipoproteins and removing excess cholesterol from sperm plasma membranes [88]. ApoA-I may also protect sperm from inflammation in the female reproductive tract [89]. Lipid metabolic processes involving apoA-I are an important feature of chicken and turkey seminal fluid [25, 26], and apoA-1 is upregulated in high-laying chicken breeds compared to wild-type [90], suggesting it may be under selection in domesticated poultry breeds due to its positive role in fertility and oviposition.

Foam-mediated calcium ion (Ca^2+^) transport and binding (GO Term: 0005509) may regulate sperm motility and the acrosome reaction. Ca^2+^ influx through ion channel proteins mediates intracellular signalling pathways and essential cell processes in mammalian sperm [91] and may mediate motility and the acrosome reaction in chicken sperm [92]. It is possible that nucelobindin 2, a highly abundant foam protein, mediates calcium ion influx through its direct interaction with the sperm midpiece, as it does in mammalian seminal plasma [93, 94].

Finally, we found evidence that foam proteins play a role in precise actin cytoskeletal organisation, essential for sperm motility, capacitation, and the acrosome reaction [95–98]. Actin cytoplasmic 2, a gamma-type actin, highly abundant in foam, may mediate motility in the sperm flagellum [99], whilst other actin filaments (Go Term: 0005884) may regulate sperm head capacitation and initiate the acrosome reaction during fertilisation [96]. Actin also functions during cytokinesis (cell cytoplasm division), vesicle and organelle movement, cell signalling, and the establishment and maintenance of cell junctions and shape [100]. The presence of these proteins suggests foam contains integral sperm cytoplasm components and regulates sperm function.

### 4.3 Role of Foam Proteins in Immune Function

Avian ejaculates are susceptible to harbouring bacterial infections because the cloaca functions in both excretion and sperm transfer. As Japanese quail are ground-dwelling, their likelihood of bacterial infection is even higher due to contact with the ground [101–103]. Bacteria can directly damage sperm form and function, reducing sperm motility and viability [104, 105], or indirectly via bacterial-induced acute inflammatory response [106]. Immunity proteins in seminal fluid can protect sperm from bacterial-induced damage [1, 107] and this may be particularly important in domesticated species where individuals often live in closer proximity than in the wild.

Immunoglobulin domain proteins in quail foam may regulate an immune response (GO Term: 0006955) in the female reproductive tract. We found in high abundance an Ig receptor (IgR) protein ortholog to the human IgR gene, which interacts with IgA antibodies in the mucosal immune system, defending against antigens [108]. Several IgR proteins are present in human seminal fluid [109], and boar (*Sus scrofa*) seminal fluid IgR proteins are associated with fewer sperm defects [110].

Foam may also defend against bacteria via an antibacterial humoral response (GO Term: 0019731). Avidin binds to biotin in bacteria to prevent growth [111], and ovoinhibitor and ovotransferrin, the two most abundant foam proteins, have antimicrobial properties which may protect sperm inside the female tract [112]. Both these proteins are found in chicken and turkey seminal fluid [24–26], and in egg albumin where they provide antimicrobial protection to embryos [113]. Ovoinhibitor acts as a serine protease inhibitor [112], whilst ovotransferrin regulates antigen genome replication [114]. Both ovoinhibitor and ovotransferrin have evolved as unique antimicrobial barriers only found in avian species.

We found innate immunity proteins in foam, which may confer immediate antimicrobial benefits. For example, lysozyme g provides defence against Gram-positive bacterium, hydrolysing peptidoglycan in bacteria [115, 116] and ameliorating oxidate stress [117]. Lysozyme g is found in seminal fluid of humans [118], fish [119], insects [117], and birds including the wild superb fairy-wren (*Malurus cyaneus*) [120], and turkeys [112, 121]. High seminal plasma lysozyme levels are associated with higher sperm velocity and motility in brown trout (*Salmo trutta*) [119], and higher sperm performance and concentration in humans [118].

In chickens, seminal fluid changes the expression of several genes expressed in the uterus that relate to immune system function, potentially suppressing the female’s immune response (although specific proteins signalling gene modulation are unknown) [122]. Artificial insemination also elicits a controlled inflammation-like response in the chicken oviduct, which causes selective sperm phagocytosis [123, 124]. In quail foam, we identified an S100 protein which regulates intracellular homeostasis and, during cellular response to stress, can be secreted extracellularly to induce a pro-inflammatory response [125–127]. This may prepare the female reproductive tract for fertilisation and embryonic development by removing pathogens and promoting immune tolerance towards sperm. We also found a predicted c-type lectin domain protein (CTL) in foam. In mammals, c-type lectins bind to carbohydrate structures on self-antigens in a calcium-dependent manner to signal tolerance [128, 129], regulating immune homeostasis and promoting sperm survival in the female reproductive tract [130, 131]. Finally, pepsins, a type of aspartic protease found in foam, are not known to affect sperm characteristics but are active in female vaginal fluid 24 hours after ejaculation in humans [132]. It is plausible that pepsins degrade seminal proteins in the female reproductive tract via proteolysis, preventing a female immunogenic response, but this remains to be tested.

### 4.4 Unique Foam Structure

We found several proteins unique to Japanese quail, without chicken orthologs. Four were predicted neuroblast differentiation-associated AHNAK-like proteins, similar to a protein that regulates calcium ion channels in chickens [133, 134]. Two unique Ig-like domain-containing proteins found in foam suggest adaptive immune function and a unique aldose reductase protein that catalyses various redox reactions may reduce oxidative stress and have immunogenic effects including carbonyl detoxification [135] and reduction of aldehyde phospholipids [136].

### 4.5 Conclusion

Our interrogation of the Japanese quail foam proteome suggests roles in sperm maturation and DNA protection, semen liquefaction, sperm plasma membrane homeostasis, and energy production for sperm motility. Our data suggest foam may play a role in protecting sperm from antigens, provide anti-inflammatory properties, and reduce/prevent immunogenic responses in the female reproductive tract to enhance fertilisation success and support embryonic development. Future experimental studies could investigate the specific function of key proteins and enzymes identified here during fertilisation dynamics and post-copulatory sexual selection. This work advances our understanding of the function of Japanese quail unique seminal foam and evolution of avian reproductive systems.

## 5 Associated Data

The mass spectrometry proteomics data have been deposited to the ProteomeXchange Consortium via the PRIDE [137] partner repository with the dataset identifier PXD063034.

## Acknowledgements

C.M. was funded by a NERC ACCE studentship (NE/S00713X/I), O.V. was funded by the German Research Foundation (DFG, Deutsche Forschungsgemeinschaft, 428800869), B.T. was funded by the Swiss National Science Foundation (PP00P3 128386 and PP00P3 157455) and N.H. was funded by a Royal Society Dorothy Hodgkin Fellowship (DHF160200). Mass spectrometry analyses were performed by the biOMICS Facility at the University of Sheffield.

